# On the origins and evolution of trans-splicing of bursicon in mosquitos

**DOI:** 10.1101/050625

**Authors:** Scott William Roy

## Abstract

Broad transcriptomic sequencing of eukaryotes has revealed the ubiquity of splicing of nuclear genes. While the vast majority of splicing events join segments of the same RNA transcript, various studies have found a few intriguing cases of trans-splicing of introns, in which splicing events within protein coding regions join segments of different RNA transcripts. The most structurally intricate case known involves the bursicon gene in mosquitos, in which an internal exon is encoded at a distinct locus, requiring multiple trans-splicing events form the mature mRNA. This arrangement is known to be ancestral to mosquitos, however the exact timing of the origin of trans-splicing and the history of the bursicon gene within mosquitos is unknown. Taking advantage of the recent availability of genomes from various *Anopheles* mosquitos and from relatives of mosquitos, I determined *trans* versus *cis* encoding of bursicon across Culicomorpha. I conclude that trans-splicing emerged in the last common ancestor of mosquitos, and that trans-splicing has been retained in all 19 studied *Anopheles* species. The retention of trans-splicing could indicate functional importance of this arrangement, or could alternatively reflect the rarity of mutations giving rise to viable allelic alternatives.

---

In eukaryotic nuclear genes, the protein-coding regions of mRNA transcripts do not represent a single contiguous genomic region, but rather are spliced together from RNA segments representing different DNA regions (Roy and Irimia 2014). Almost always, these segments represent subsegments of a single RNA transcript which are joined by a molecular machine called the spliceosome. Portions that are retained in the mature mRNA are termed exons while the internal removed sections are termed introns. Much more rarely, the protein-coding region of an mRNA can be formed by splicing together portions of RNA transcripts transcribed from different genomic loci, that is, trans-splicing of introns (Li et al. 1999; Takahara et al. 2000; Dorn et al. 2001; Robertson et al. 2007; Nageshan et al. 2011; Kamikawa et al. 2011; Roy et al. 2012). (This project should be distinguished from so-called spliced leader trans-splicing, in which a short RNA tag (the spliced leader) is added to the 5 end of various mRNA species, typically donating a (partial) UTR (Lasda and Blumenthal 2011)). Trans-splicing of introns within coding regions is thought to be very uncommon, with only a handful of cases known across eukaryotes.

The most structural intricate known case of nuclear trans-splicing involves bursicon, an important developmental hormone in insects (Robertson et al. 2007). In characterized mosquitos, the mature bursicon mRNA consists of four exons, derived from two pre-mRNA transcripts transcribed from different genomic loci. One transcript encodes exons 1, 2 and 4, and the second encodes exon 3. Two spliceosomal reactions occur between the two RNA species, one joining the 3 end of exon 2 to the 5 end of exon 3 and the other joining the 3 end of exon 3 to the 5 end of exon 4 (Figure 1a). This arrangement is known to be ancestral to mosquitos as it is found in the divergent species *Anopheles gambiae, Aedes aegypti* and *Culex pipiens* (Robertson et al. 2007). However, the timing of the origin of this arrangement, and its evolutionary history within other lineages of mosquitos, remain obscure.

**Figure 1.**
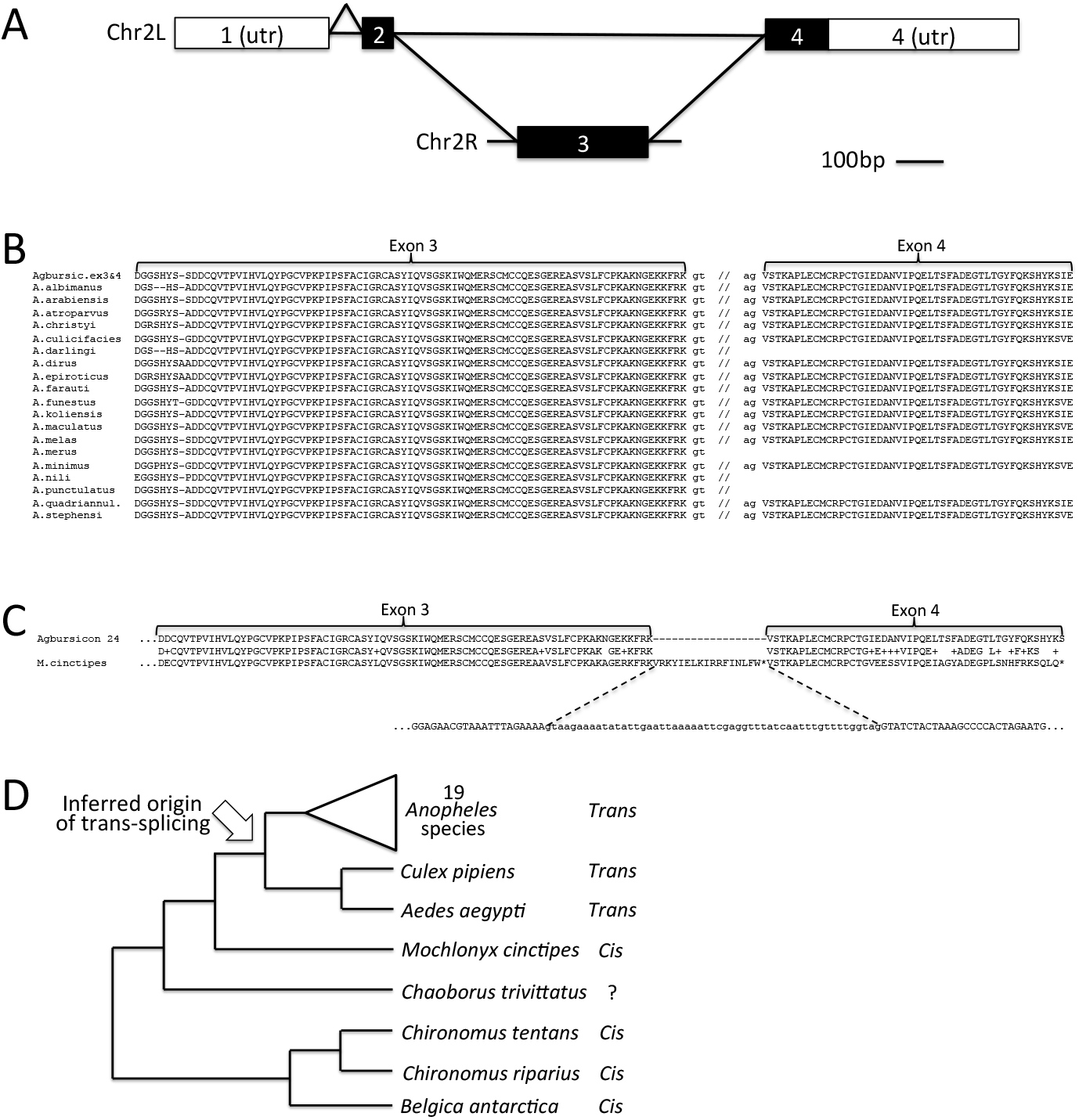
Evolutionary history of the trans-spliced bursicon gene in mosquitos. A. Schematic of expression of the bursicon gene in *A. gambiae*, adapted from Robertson [2007]. Two pre-mRNA transcripts are produced, one from chromosome 2L and one from chromosome 2R. The former contains three exons (1,2 and 4, with exon 1 encoding the 5′ UTR and exon 4 encoding the C-terminus of the protein and the 3′ UTR], and the latter contains one exon (exon 3]. The mature message is created by three splicing reactions including two joining portions of the two pre-mRNA transcripts. B. Alignment of the *A. gambiae* bursicon protein with homologous sequences for the 19 species of *Anopheles* mosquito characterized here. For each species for which sequence homologous to both exon 3 and exon 4 were found, they were found on different DNA molecules, suggesting trans-splicing. C. Alignment of the *A. gambiae* bursicon protein with homologous sequence on contig JXPH01093764.1 in the *M. cinctipes* genome, showing *cis* orientation of exons 3 and 4 and an intervening putative intron with gt…ag splice boundaries and in-frame stop codons. D. Phylogeny of studied species within the infraorder Culicomorpha, showing species with evidence for *trans* or *cis* encoding of bursicon.

To trace the history of the bursicon gene, I downloaded recently available genomes from various mosquitos and related species (Vicoso and Bachtrog 2015; Neafsey et al. 2015). First, I downloaded species within the very ancient genus *Anopheles* (dating back ~100My; see Neafsey et al. 2015) and performed TBLASTN searches of the *A. gambiae* bursicon protein sequence against each species genome. For all 15/19 searched species, sequences with very high levels of protein-level similarity to exons 3 and 4 of *A.gambiae* bursicon were found on different DNA molecules, consistent with a pattern of trans-splicing similar to that found in previously characterized mosquitos. For the other 4 species, a single hit was found to exon 3, but no sequence (either proximal or distal to the exon 3 hit) was found for exon 4. One possible explanation for this pattern is that the exon 4 sequence is not represented in the current genome assembly. For no species did I find a sequence homologous to exon 2, possibly because exon 2s function in encoding a signaling peptide gives it a higher degree of protein sequence flexibility (and thus rate of evolution). All these results are consistent with retention of the *A.gambiae* trans-splicing pattern across *Anopheles* species.

To trace the origins of the trans-splicing arrangement in mosquitos, I downloaded the genomes of five related species within the larger infraorder Culicomorpha. TBLASTN searches of the *A*gambiae bursicon protein gave clear hits for four of the five genomes. Figure 1. shows the phylogeny of the studied species. For all cases, sequences homologous to exon 3 and exon 4 appeared on the same genomic DNA molecule in the order expected for co-linear transcription within a single molecule. For all cases, intervening sequences between the exon 3 and exon 4 regions contained stop codons or frameshifts, strongly suggesting the presence of a *cis*-spliced intron at or near the position of the trans-spliced intron in mosquitos. This suggests that trans-splicing evolved in the ancestor of mosquitos.

## Concluding remarks

It is striking that the peculiar arrangement of the trans-spliced bursicon gene has been independently retained in a wide variety of mosquitos. Conservation is often taken as evidence of function, and the possibility that the trans-spliced arrangement imparts a function that could not be accomplished by a cis-spliced form is enticing. However, an important alternative is that production of viable cis-spliced alternatives by random mutation is an extremely rare occurrence. Reinsertion of exon 3 into the genomic region between exons 2 and 4 would require a very precise non-homologous double recombination, and even then might lead to mis-splicing due to disruption of splicing signals or to disrupted functioning of splicing signals in cis. Another possibility is retroposition of a spliced mRNA at a third genomic locus, however such a gene would likely lack various intronic and upstream regulatory motifs, and thus might not be able to replace the ancestral function.

Additional data. Results from TBLASTN search of *A gambiae* bursicon protein against various *Anopheles* species genomes. For each line, the format is subject_scaffold:genomic_coordinates, then the amino acid positions of the *A gambiae* bursicon protein that fall within the BLAST hit.

A.albimanus

KB672320.1:524518-524784 Agbursicon:25-115

KB672342.1:143410-143261 Agbursicon:114-163

::::::::::::::

A.arabiensis

KB704451.1:4928193-4928459 Agbursicon:27-115

KB704396.1:2542421-2542570 Agbursicon:114-163

::::::::::::::

A.atroparvus

AXCP01008157.1:7448-7170 Agbursicon:23-115

KI421891.1:6686038-6685889 Agbursicon:114-163

::::::::::::::

A.christyi

KB697805.1:644-922 Agbursicon:23-115

KB678907.1:3115-3285 Agbursicon:107-163

::::::::::::::

A.culicifacies

AXCM01005500.1:1165-893 Agbursicon:27-117

AXCM01017223.1:760-611 Agbursicon:114-163

::::::::::::::

A.darlingi

ADMH02002125.1:942117-941851 Agbursicon:27-117

::::::::::::::

A.dirus

KB672979.1:6926299-6926015 Agbursicon:28-121

KB672602.1:5830665-5830513 Agbursicon:113-163

::::::::::::::

A.epiroticus

KB671576.1:75830-75540 Agbursicon:20-115

KB671690.1:102779-102928 Agbursicon:114-163

::::::::::::::

A.farauti

KI915040.1:593704-593438 Agbursicon:27-115

KI915044.1:11246457-11246606 Agbursicon:114-163

::::::::::::::

A.funestus

KB669425.1:142513-142241 Agbursicon:27-117

KB669203.1:467130-466981 Agbursicon:114-163

::::::::::::::

A.koliensis

JXXB01041064.1:3306-3025 Agbursicon:22-115

JXXB01016478.1:5058-4909 Agbursicon:114-163

::::::::::::::

A.maculatus

AXCL01025102.1:272-6 Agbursicon:27-115

AXCL01011182.1:483-319 Agbursicon:109-163

::::::::::::::

A.melas

AXCO02002142.1:785-1087 Agbursicon:27-127

KI919430.1:57035-56874 Agbursicon:110-163

::::::::::::::

A.merus

KI915301.1:220765-220499 Agbursicon:27-115

KI915256.1:770693-770532 Agbursicon:110-163

::::::::::::::

A.minimus

KB663610.1:4819334-4819062 Agbursicon:27-117

KB663655.1:917356-917207 Agbursicon:114-163

::::::::::::::

A.nili

ATLZ01013352.1:519-253 Agbursicon:27-115

::::::::::::::

A.punctulatus

JXXA01016707.1:1714-1980 Agbursicon:27-115

::::::::::::::

A.quadriannulatus

KB667777.1:4536606-4536878 Agbursicon:27-117

KB668088.1:1031074-1030925 Agbursicon:114-163

::::::::::::::

A.sinensis

KE524456.1:61412-61708 Agbursicon:17-115

KE525275.1:769221-769370 Agbursicon:114-163

::::::::::::::

A.stephensi

KE388949.1:917632-917354 Agbursicon:23-115

ALPR02015437.1:909-760 Agbursicon:114-163

